# A *Citrullus* genus super-pangenome reveals extensive variations in wild and cultivated watermelons and sheds light on watermelon evolution and domestication

**DOI:** 10.1101/2023.06.08.544282

**Authors:** Shan Wu, Honghe Sun, Lei Gao, Sandra Branham, Cecilia McGregor, Susanne S. Renner, Yong Xu, Chandrasekar Kousik, W. Patrick Wechter, Amnon Levi, Zhangjun Fei

## Abstract

Sweet watermelon (*Citrullus lanatus* subsp. *vulgaris*) is among the most important vegetable crops in the world. Wild relatives are important resources for watermelon breeding. Here we report high-quality reference genomes of three wild watermelons, *C. mucosospermus, C. amarus* and *C. colocynthis*, and the divergence and genome evolution of different *Citrullus* species. Using genomic data from 547 watermelon accessions spanning four *Citrullus* species, we construct a super-pangenome to represent the *Citrullus* gene repertoire and provide a catalogue of orthologous relationships among species. Gene presence/absence variation analysis uncovers many disease resistance genes that are missing in cultivated watermelons, as well as genes with significantly different occurrence frequencies between populations that might underlie watermelon evolution and domestication. We revisit watermelon domestication using the recently identified wild progenitor, Kordofan melon, which provides insights into the domestication of fruit bitterness, sweetness and flesh coloration. The *Citrullus* super-pangenome provides a valuable resource for breeding and biological discovery, and our comparative genomic analyses shed additional light on watermelon evolution and domestication.

## Introduction

Watermelon (*Citrullus lanatus* subsp. *vulgaris*) has been domesticated for at least 4000 years (Paris, 2015; Renner et al., 2021). It is among the most important vegetable crops widely grown in temperate and tropical regions of the world, with over 3 million hectares planted and the production exceeding 101 million tons in 2020 (https://www.fao.org/faostat). Cultivated watermelon suffers from various diseases and abiotic stresses, and has a narrow genetic base due to domestication and breeding focusing primarily on fruit quality traits (Levi et al., 2001), which has challenged the improvement of watermelon for disease resistance. Bitter or bland-tasting wild watermelons, such as *C. mucosospermus, C. amarus* and *C. colocynthis*, are valuable sources of resistance to many diseases including Fusarium wilt, powdery mildew, Phytophthora fruit rot, gummy stem blight, anthracnose and various viruses (Levi et al., 2017). Weak reproductive barriers among most species in the *Citrullus* genus have allowed the usage of wild watermelons in genetic studies and the improvement of modern cultivars (Guo et al., 2019).

Recently, a wild form of watermelon from Sudan, known as the Kordofan melon (*C. lanatus* subsp. *cordophanus*), was identified as the closest relative and a possible direct progenitor of domesticated watermelon (Renner et al., 2021). The Kordofan melon carries non-bitter fruits and shares a much more similar genetic background with domesticated watermelon than any other extant wild watermelons. Using Kordofan melons as breeding materials could bring fruit quality and/or disease resistance-related genetic elements that have been lost during domestication back to modern cultivars, while avoiding the introduction of many undesirable traits such as those present in more distantly related wild watermelons. However, the potential of Kordofan melon in providing beneficial traits for modern cultivars has been largely unexplored, and this wild material has not been utilized in the study of watermelon domestication and in any watermelon breeding programs.

Three *C. lanatus* reference genomes have been released (Guo et al., 2019; Wu et al., 2019; Renner et al., 2021), which have greatly facilitated the discovery of genetic loci associated with agronomically important traits including disease resistance. Enabled by a recent surge in plant genome sequencing, pan-genomes that represent the genetic diversity within species have greatly supported plant breeding and evolution studies (Bayer et al., 2020; Della Coletta et al., 2021). Compared with pan-genomes at the species level, super-pangenomes at the genus level provide a more complete genomic variation repertoire covering useful genetic elements underlying beneficial traits lost during domestication but preserved in related wild species (Khan et al., 2020). For example, including both cultivated and wild rice species in a super-pangenome study has deepened our understanding of rice adaptation and domestication (Shang et al., 2022). A *Citrullus* genus super-pangenome comprising all genetic elements from cultivated watermelon and its wild relatives is not only crucial for studying molecular mechanisms underlying watermelon traits, but also provides an insightful resource for exploring the evolution and domestication histories of watermelon, and for more efficient utilization of beneficial genes and alleles in watermelon breeding.

In this study, we assembled high-quality genomes for three wild watermelons, *C. mucosospermus* USVL531-MDR, *C. amarus* USVL246-FR2, and *C. colocynthis* PI 537277. The reference genomes were used for comparative genomic and phylogenetic analyses to study the divergence times and chromosome karyotype evolution of watermelons. We further performed genome resequencing of 201 wild watermelon accessions from *C. colocynthis, C. amarus, C. mucosospermus*, and *C. lanatus* subsp. *cordophanus*. We constructed four species-level watermelon pan-genomes and a *Citrullus* super-pangenome from six reference-grade genome assemblies and resequencing data of 541 accessions, highlighting the diverse genetic background of different watermelon species and delivering a catalogue of orthologous gene relationships between cultivated watermelon and its wild relatives. Gene presence/absence variation (PAV) analyses revealed functionally important genes that could be selected during watermelon evolution and domestication. Finally, we revisited the history of watermelon domestication by using the Kordofan melon as the progenitor to identify selective sweeps and selected genes controlling fruit quality traits.

## Results

### Reference genome assemblies of *C. mucosospermus, C. amarus* and *C. colocynthis*

Three wild watermelon accessions were selected for reference genome sequencing: *C. mucosospermus* USVL531-MDR, resistant to powdery mildew and Phytophthora fruit rot (Mandal et al., 2020); *C. amarus* USVL246-FR2, resistant to Fusarium wilt (Wechter et al., 2016) and bacterial fruit blotch (Branham et al., 2010); and *C. colocynthis* PI 537277, resistant to whiteflies (Coffey et al., 2015) and Papaya ringspot virus-watermelon strain (Levi et al. 2016). Genome sizes of USVL531-MDR, USVL246-FR2 and PI 537277 were estimated to be 434.7 Mb, 423.2 Mb and 406.0 Mb, respectively, based on k-mer depth distribution analyses of Illumina sequences (**Supplementary Fig. 1**). For USVL531-MDR, we generated 40.1 Gb PacBio sequences, covering 92.3× of the genome (**Supplementary Table 1**). The PacBio reads were assembled into contigs followed by polishing with PacBio and Illumina reads, which resulted in an assembly containing 77 contigs with a total size of 365.3 Mb and an N50 length of 27.6 Mb (**Supplementary Table 2**). Using the synteny to the cultivar 97103 reference genome, 99.36% of the USVL531-MDR assembly (363.0 Mb; 21 contigs) were anchored to 11 chromosomes (**Supplementary Table 3**). Ten out of the 11 USVL531-MDR chromosomes were each composed of only one or two contigs. The genomes of *C. amarus* USVL246-FR2 and *C. colocynthis* PI 537277 were sequenced using the Illumina technology. The high-quality cleaned sequences produced from paired-end and mate-pair libraries covered approximately 284.2× and 370.1× of the USVL246-FR2 and PI 537277 genomes, respectively (**Supplementary Table 1**). The genome assembly of USVL246-FR2 had a total size of 386.7 Mb, and consisted of 38,258 contigs and 1,422 scaffolds, with N50 sizes of 18.1 kb and 3.8 Mb, respectively (**Supplementary Table 2**). Using genetic maps generated from the recombinant inbred line (RIL) and F2 populations derived from a cross between USVL246-FR2 and USVL114 (Branham et al., 2019), 95.9% of the assembly was anchored to 11 linkage groups and 93.6% of the assembly was oriented (**Supplementary Fig. 2** and **Supplementary Table 3**). The PI 537277 genome was assembled into 15,928 contigs and 1,536 scaffolds with N50 sizes of 44.9 kb and 1.6 Mb, respectively, and the total assembly size was 356.1 Mb. About 99.7% of the PI 537277 assembled sequences were constructed into 11 pseudochromosomes using Hi-C data (**Supplementary Fig. 3** and **Supplementary Table 3**). Completeness of the three assemblies was evaluated by BUSCO (Simao et al., 2015), which revealed that 99.07%, 98.95% and 99.01% of the core conserved plant genes were detected complete in the USVL531-MDR, USVL246-FR2 and PI 537277 assemblies, respectively (**Supplementary Table 4**). Aligning genomic reads back to the assembly revealed mapping rates of 98.58%, 98.72% and 99.67% for USVL531-MDR, USVL246-FR2 and PI 537277, respectively. The LAI values (Ou et al., 2018) were 10.66, 7.17 and 8.53 for USVL531-MDR, USVL246-FR2 and PI 537277, respectively, comparable to that of the high-quality 97103 reference genome (9.96; Guo et al., 2019). Collectively, these results demonstrated the high quality of the three assemblies.

### Gene contents and divergence among watermelon species

The repeat contents in the assembled *C. mucosospermus* USVL531-MDR, *C. amarus* USVL246-FR2 and *C. colocynthis* PI 537277 genomes were 45.28% (165.4 Mb), 42.83% (166.2 Mb) and 48.07% (175.6 Mb), respectively (**Supplementary Table 5**). Each repeat-masked assembly was annotated for protein-coding genes using a combination of *ab initio* gene prediction coupled with evidence from transcript mapping and protein homology. To further improve gene predictions and reduce the impact of annotation difference on false discovery of gene presence/absence across species, genes were mapped with the Liftoff software (Shumate and Salzberg, 2021) among the six watermelon reference genomes including three reported in this study and three published ones, *C. lanatus* 97103 (Guo et al., 2019), *C. lanatus* Charleston Gray (Wu et al., 2019) and Kordofan melon (*C. lanatus* subsp. *cordophanus*) (Renner et al., 2021). A total of 21,676 to 22,723 protein-coding genes were predicted in these six genomes of different watermelon species or subspecies (**Supplementary Table 4**). BUSCO assessment showed that 94.55% to 96.01% of the plant core genes were captured complete in genes predicted from the six watermelon genomes (**Supplementary Table 4**). A total of 1,516, 1,591 and 1,092 additional genes were annotated in the published reference genomes of 97103, Charleston Gray and Kordofan melon, respectively, and 10,121, 8,744 and 9,393 previous gene models were updated.

Molecular dating using single-copy orthologous genes in genomes of the five watermelon species and subspecies, seven other cucurbit species, strawberry and walnut revealed that *Citrullus* species diverged from their sister clade 13.79 (10.30 to 17.39) million years ago (Mya) (**Fig. 1a**), consistent with the previous finding (Wu et al., 2017). The divergence between cultivated watermelon and wild Kordofan melon was dated to be 0.18 (0.12 to 0.24) Mya, and *C. lanatus* split from its West African sister, *C. mucosospermus*, 0.25 (0.17 to 0.32) Mya. The inferred divergence time between cultivated watermelon and its direct progenitor, Kordofan melon, was earlier than the anticipated time of watermelon domestication (Renner et al., 2021), and could be an overestimation caused by genes exhibiting incomplete lineage sorting due to the closely timed divergence events among *C. mucosospermus* and the two *C. lanatus* subspecies. Genome alignments between wild and cultivated watermelons revealed one-to-one chromosome-level syntenic relationships among *C. lanatus, C. mucosospermus* and *C. amarus*, with several large inversions (**Supplementary Fig. 4**). A large inter-chromosomal rearrangement involving chromosomes 1 and 4 was found between *C. colocynthis* and the other three *Citrullus* species, *C. lanatus, C. amarus* and *C. mucosospermus* (**Fig. 1b** and **Supplementary Fig. 4**). Comparative analysis of genomes of watermelons and melon (Castanera et al., 2020) revealed a high level of collinearity between the entire chromosome 4 of *C. colocynthis* and chromosome 8 of melon (**Fig. 1b**), suggesting that *C. colocynthis* likely carried the ancestral karyotype and that the inferred chromosome fission and fusion events occurred approximately 4.54-2.41 Mya after the divergence of *C. colocynthis* from other watermelons and before the separation of *C. amarus* from *C. mucosospermus* and *C. lanatus* (**Fig. 1b**).

**Figure 1.**
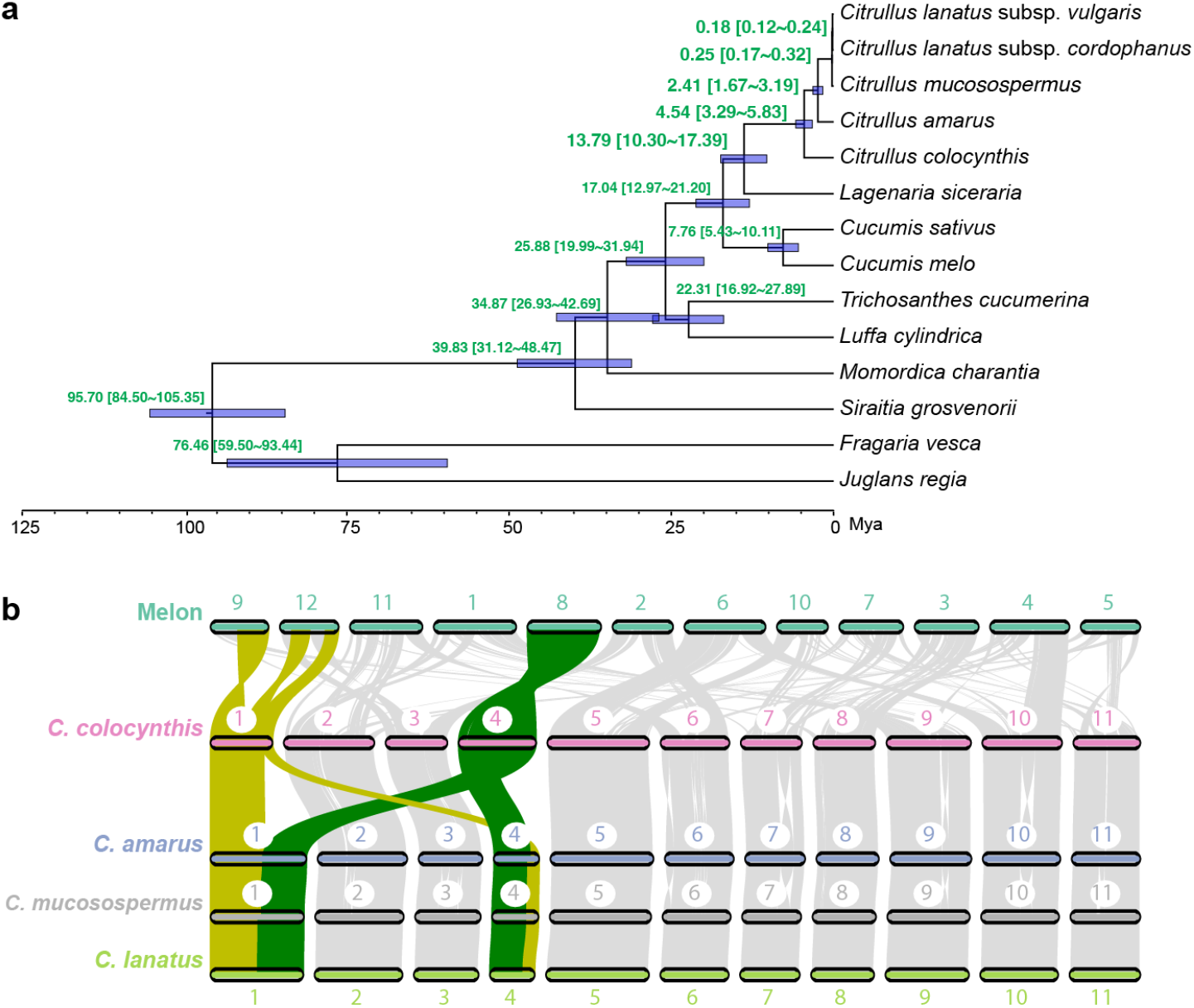
Divergence and genome evolution of watermelon species. (**a**) Phylogenetic tree and estimated times of divergence events. Green numbers near each node indicate the estimated divergence time in millions of years and the range of the 95% highest probability density intervals. Horizontal blue bar on each node indicates the range of the estimated divergence time. Genomes of five watermelons (*Citrullus lanatus* subsp. *vulgaris, C. lanatus* subsp. *cordophanus, C. mucosospermus, C. amarus* and *C. colocynthis*), bottle gourd (*Lagenaria siceraria*), cucumber (*Cucumis sativus*), melon (*Cucumis melo*), snake gourd (*Trichosanthes cucumerina*), sponge gourd (*Luffa cylindrica*), bitter gourd (*Momordica charantia*), monk fruit (*Siraitia grosvenorii*), woodland strawberry (*Fragaria vesca*) and walnut (*Juglans regia*) were used in the analysis. (**b**) Synteny among the genomes of melon, *C. colocynthis, C. amarus, C. mucosospermus* and *C. lanatus*. Genomic regions syntenic to *C. colocynthis* chromosomes 1 and 4 are highlighted in yellow and green, respectively.

### A super-pangenome of watermelon

In watermelon breeding, useful genes can be transferred from wild species to cultivars by crossing. A *Citrullus* genus-level super-pangenome has the potential to represent the complete gene repertoire of watermelons that can be utilized in current breeding programs. In this study, we resequenced 201 wild watermelon accessions, including 15 *C. lanatus* subsp. *cordophanus*, 30 *C. mucosospermus*, 122 *C. amarus*, and 34 *C. colocynthis*, in addition to one that was later identified to be a landrace. Combined with those selected for reference genomes and the ones with previously generated resequencing data (Guo et al., 2019), a total of 547 watermelon accessions were used in the watermelon super-pangenome construction, including 349 *C. lanatus* (243 cultivars, 88 landraces and 18 *C. lanatus* subsp. *cordophanus*), 31 *C. mucosospermus*, 131 *C. amarus* and 36 *C. colocynthis* (**Supplementary Table 6**). The newly generated data in this study increased the sequencing depth and number of wild accessions, especially for *C. amarus* and *C. colocynthis*. Considering the divergence between cultivated and wild watermelons, species-level pan-genomes were first constructed and then combined into a *Citrullus* super-pangenome to reduce the bias of sequence mapping between distantly related species and to distinguish novel sequences from divergent orthologs. Four species-level pan-genomes were developed using an assembly and mapping approach (**Supplementary Fig. 5a**), each containing the species-specific reference sequences, and a total of 24.5 Mb, 15.6 Mb, 18.3 Mb and 42.4 Mb non-redundant novel sequences for *C. lanatus, C. mucosospermus, C. amarus* and *C. colocynthis*, respectively, harboring 2,288, 583, 1,922 and 2,521 novel genes that were not present in the species-specific reference genomes (**Supplementary Table 7**).

To combine the four species-level pan-genomes, genes in each pan-genome were mapped to the other three pan-genomes to establish orthologous relationships between genes from difference species (gene-to-gene), and between aligned genes and genomic regions without predicted genes (gene-to-location) (**Supplementary Fig. 5b**). As a result, 34,910 orthologous groups, including 33,697 syntenic orthologous groups, 1,166 orthologous groups without syntenic information, and 47 species-specific groups, as well as 3,145 singletons, were identified (**Supplementary Table 8**). The number of genes in the *Citrullus* super-pangenome (38,055 including orthologous groups and singletons) was about 58% larger than that of the predicted genes in each species (24,090 on average), mainly because of the inclusion of gene-to-location orthologous pairs. When keeping genomic regions without annotated genes but aligned by genes from other species, 28,607 (75.2%) orthologous groups contained genes or sequences from all four species (**Fig. 2a**), among which 27,438 (95.9%) had one gene or one location in each species (1:1:1:1). However, when only considering gene-to-gene orthologous relationships, the proportion of species-specific genes increased from 8.4% (3,193 including species-specific groups and singletons) to 42.6% (16,217) (**Supplementary Fig. 6a**), with *C. colocynthis* having the most species-specific genes (**Supplementary Fig. 6b**), consistent with its most distant phylogenetic relationship to the other three watermelon species. These results together suggested that during watermelon evolution, mutations in genes such as those causing premature stops or frame shifts had occurred and possibly led to disruptions of gene functions as indicated by the absence of predicted gene models in those orthologous genomic regions; however, the sequences derived from the ancestral genes were largely maintained in the genomes rather than being completely purged. Genes that are unique to *C. lanatus* might include the ones underlying fruit quality traits that distinguish it from its wild relatives. There were 701 *C. lanatus*-specific orthologous groups and another 627 groups contained *C. lanatus*-specific tandem duplicated copies (at least two copies of *C. lanatus* genes in the syntenic region, while at most one was found in each of the other three species) (**Supplementary Table 9**). These genes included the tonoplast sugar transporter gene, *ClTST2* (*Cla97C02G036390* and *Cla97C00G000440*), which functions in regulating sugar accumulation in watermelon flesh (Ren et al., 2018). This *ClTST2* tandem duplication has also been found in the elite watermelon line G42 (Deng et al., 2022).

**Figure 2.**
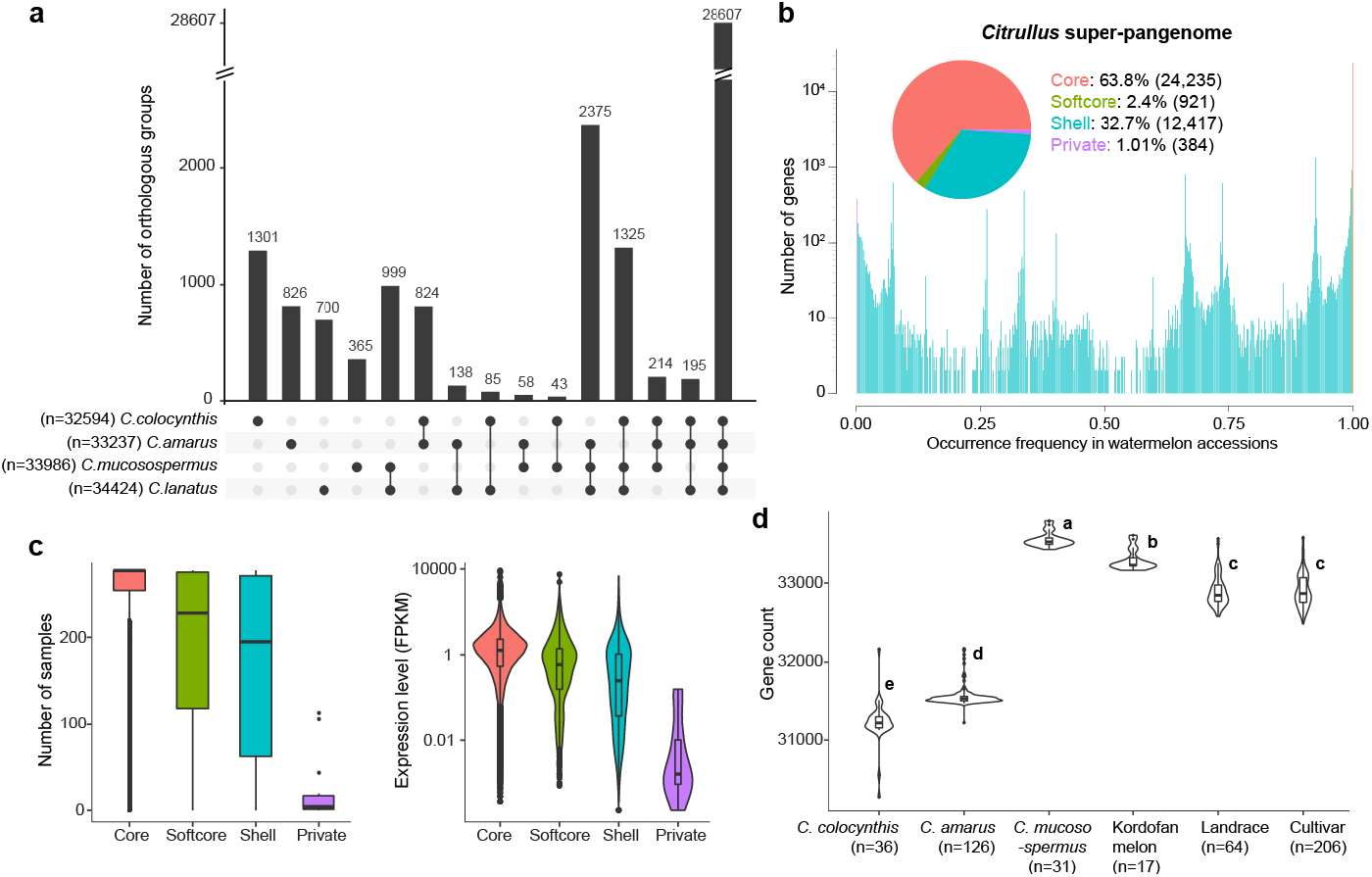
*Citrullus* super-pangenome. (**a**) Upset diagram of orthologous groups among the four watermelon species. (**b**) Composition of the *Citrullus* super-pangenome. (**c**) Expression of different categories of genes in the *Citrullus* super-pangenome derived from publicly available watermelon RNA-Seq datasets archived in CuGenDBv2. The number of samples in which the genes were expressed (left) and the average gene expression levels (right) are shown. (**d**) Numbers of genes (including orthologous loci without annotated genes) detected in individuals from different watermelon populations. Different letters in violin plots indicate significant differences among groups evaluated by Tukey’s HSD test (α<0.05).

Resequencing reads of each watermelon accession were aligned to the pan-genome of its own species for detecting gene PAVs. To control false calls of gene PAVs, accessions with insufficient read coverage were excluded, leaving 480 accessions (64 landraces, 206 modern cultivars, 17 *C. lanatus* subsp. *cordophanus*, 31 *C. mucosospermus*, 126 *C. amarus* and 36 *C. colocynthis*) for the investigation of gene PAVs (**Supplementary Table 6**). Genes in the pan-genome were classified as core (present in all accessions), softcore (present in all but one), private (present in only one accession), and shell (between softcore and private). In the *Citrullus* super-pangenome, the core gene content was 63.8% (24,235 genes) (**Fig. 2b**), much lower than those in the species-level pan-genomes (85.6%, 97.2%, 90.0% and 88.7% for *C. lanatus, C. mucosospermus, C. amarus* and *C. colocynthis*, respectively) (**Supplementary Fig. 7**), indicating diverse genetic makeups among the four species. Compared to genes in the dispensable portion, core genes were expressed at higher abundance and in more watermelon samples (**Fig. 2c**), further supporting that shell and private genes were more dispensable. The numbers of orthologous genes and genomic locations found in the four species ranged from 32,594 to 34,425. *C. colocynthis* and *C. amarus* individuals carried significantly fewer genes/loci than *C. mucosospermus* and *C. lanatus* individuals (**Fig. 2d**). However, regarding the predicted genes, *C. colocynthis* had the most genes both at the species level and in individual genomes (**Supplementary Fig. 6c**). These results revealed differences in the loss of ancestral gene loci and the retention of functional genes in the genomes of various *Citrullus* species, likely due to lineage-specific evolutionary events. Fewer genes/loci were found in genomes of cultivated individuals than in those of Kordofan melons (**Fig. 2d** and **Supplementary Fig. 6c**), suggesting gene loss during watermelon domestication, a phenomenon that has also been reported in tomato (Gao et al., 2019).

### Selection of genes during watermelon evolution and domestication

Genes with different occurrence frequencies between watermelon species or groups include those involved in adaptation and/or domestication. Pairwise comparisons among the four species were carried out and genes with significantly different presence frequencies (FDR < 0.001 and fold change > 2) between species were identified (**Supplementary Tables 10** and **11**). About five to seven thousand such genes were found between any two of the four *Citrullus* species, except that only about one thousand genes with significantly altered frequencies were found between the closely related *C. lanatus* and *C. mucosospermus*. For all species, when compared with another species, genes with significantly increased frequencies were enriched with biological processes related to ‘photosynthesis’ and ‘nucleotide metabolic process’ (**Supplementary Table 12**). These results suggest that after speciation, different watermelons had kept different sets of genes participating in these most basic biological processes in plants. This might be a result of random losses of functionally redundant genes or represent a fine-tuning of these pathways through actively retaining specific genes in response to the environmental conditions that different watermelon species were exposed to. Interestingly, in comparison to *C. colocynthis* and *C. amarus*, genes with increased frequencies in *C. mucosospermus* and *C. lanatus* were enriched with biological processes ‘meristem maintenance’ and ‘meristem development’. Of the 24 genes associated with these biological processes and with significantly higher frequencies in *C. mucosospermus* and *C. lanatus*, 18 encoded serine/threonine-protein phosphatase 7 long form proteins (**Supplementary Table 13**), orthologous to Arabidopsis *MAINTENANCE OF MERISTEMS LIKE 3* that is expressed in the vasculature and hydathodes of leaves (Ühlken et al., 2014). PAV of these genes may contribute to differences in leaf growth observed among watermelon species (Levi et al., 2017).

We compared *C. lanatus* subsp. *vulgaris* (including landraces and cultivars) to *C. colocynthis, C. amarus* and *C. mucosospermus*, three wild species that have been found to be resistant to various watermelon diseases, and identified 17 genes related to disease resistance that have been lost or are present at low frequencies in the *C. lanatus* gene pool (**Supplementary Table 14**). Among these genes, five were present in multiple wild species, and five were specific to *C. colocynthis*, five to *C. amarus*, and two to *C. mucosospermus*. Most of these disease resistance genes from the wild watermelons were present in most accessions of the species, suggesting that they could be important for the survival and thrive of the species. There were also 17 disease resistance genes with significantly increased occurrence frequencies in *C. lanatus* in comparison to *C. colocynthis* and/or *C. amarus* (**Supplementary Table 14**). These genes were present in almost all *C. lanatus* and *C. mucosospermus* accessions, suggesting lineage-specific retention. By comparing *C. lanatus* landrace to its progenitor, Kordofan melon, 145 genes were found to have significantly changed occurrence frequencies during domestication (130 and 15 with decreased and increased frequencies, respectively), while during improvement from landraces to cultivars, only 13 genes had significantly altered frequencies (11 decreased and 2 increased) (**Supplementary Tables 10** and **15**).

Fruit quality traits including flesh sweetness and coloration changed dramatically during watermelon domestication and improvement. Our results suggested that these changes might be achieved mainly through mutations modifying the function of existing genes or their expression patterns, rather than removal or introduction of entire genes. A gene encoding late embryogenesis abundant protein, *Cla97C06G117170*, had a decreased frequency in cultivated watermelon compared to Kordofan melon and co-located with QTLs controlling seed traits (**Supplementary Table 15**). The potential function of *Cla97C06G117170* suggested its role in seed embryo development. This gene could be crucial for the survival of watermelons in the wild but might have become less important under cultivated conditions. There were other genes found in QTLs and/or abundantly expressed in developing fruits (**Supplementary Table 15**), yet their functions require further investigation.

### Revisiting the domestication of watermelon

We previously used *C. mucosospermus* to represent the wild progenitor of cultivated watermelon for domestication sweep detection, due to its close phylogenetic relationship to *C. lanatus* and its primitive fruit flesh characteristics, such as the lack of both sweetness and coloration (Guo et al., 2019). In this study, we included 18 lines of Kordofan melon, a descendant of the possible direct progenitor of cultivated watermelon (Renner et al., 2021). A total of 13,256,154 and 2,277,760 high-quality SNPs and small insertions/deletions (indels) were identified among the 547 watermelon accessions. Among the SNPs, 260,380 led to non-synonymous changes and 11,066 led to start/stop codon gain/loss, and 28,445 small indels caused coding sequence changes (**Supplementary Table 16**). Phylogenetic analysis using SNPs at fourfold degenerate sites revealed that the 18 Kordofan melons formed a monophyletic clade that was more closely related to cultivated watermelons than any other wild species (**Supplementary Fig. 8**). The nucleotide diversity (π) in Kordofan melons (n = 18) was 0.84×10^−3^, higher than that of landraces (n = 88; π = 0.71×10^−3^) and cultivars (n = 243; π = 0.58×10^−3^). These results together supported the current hypothesis that Kordofan melon in Northeast Africa is the progenitor of the cultivated watermelon rather than a feral form (Renner et al., 2021).

To study the watermelon genomic landscape altered by domestication, we compared landrace to Kordofan melon, and identified 123 domestication sweeps with a cumulative length of 17.62 Mb (**Supplementary Table 17**) and harboring 399 annotated genes, among which 107 were in fruit quality QTLs, including eight controlling flesh sweetness on chromosomes 2, 3 and 8, two controlling fruit weight on chromosomes 2 and 3, and five controlling fruit shape on chromosomes 2, 3, 4 and 10 (**Fig. 3a** and **Supplementary Table 18**). One of these genes, *Cla97C05G101010*, encoding a cytochrome P450 CYP82D47-like protein, was not identified previously in domestication sweeps using *C. mucosospermus* as the progenitor (Guo et al., 2019), and was in a rind thickness QTL, *Qrth5* (Sandlin et al., 2012). Its high expression levels in the fruit rind of watermelon 97103 (**Supplementary Fig. 9**; Guo et al., 2013) made it a strong candidate gene for this trait. Bitterness of fruit flesh is variable in *C. mucosospermus*, and the non-bitter trait has been fixed in cultivated watermelon (Guo et al., 2019). None of the Kordofan melons were bitter and all of them carried the homozygous non-bitterness allele of the *ClBt* gene (**Supplementary Table 19**), which contains a mutation leading to a premature stop codon and loss-of-function of the gene (Zhou et al., 2016). In contrast to *C. mucosospermus*, the genomic region surrounding *ClBt* had a low genetic diversity in Kordofan melon comparable to that in cultivated watermelon (**Supplementary Fig. 10a**), suggesting that loss of bitterness in fruit flesh had already happened in the progenitor of cultivated watermelon prior to domestication. Kordofan melons displayed white or pinkish fruit flesh colors (**Supplementary Fig. 11**) and carried different alleles at the SNP site in the *lycopene β-cyclase* (*LCYB*) gene that controls fruit flesh color in watermelon (**Supplementary Table 19**), and the SNP leads to an amino acid change from a conserved phenylalanine to valine (Bang et al., 2007). Kordofan melons had kept higher genetic diversity in the *LCYB* genomic region in comparison to landraces and cultivars (**Supplementary Fig. 10b**), suggesting that *LCYB* has been selected during watermelon domestication.

**Figure 3.**
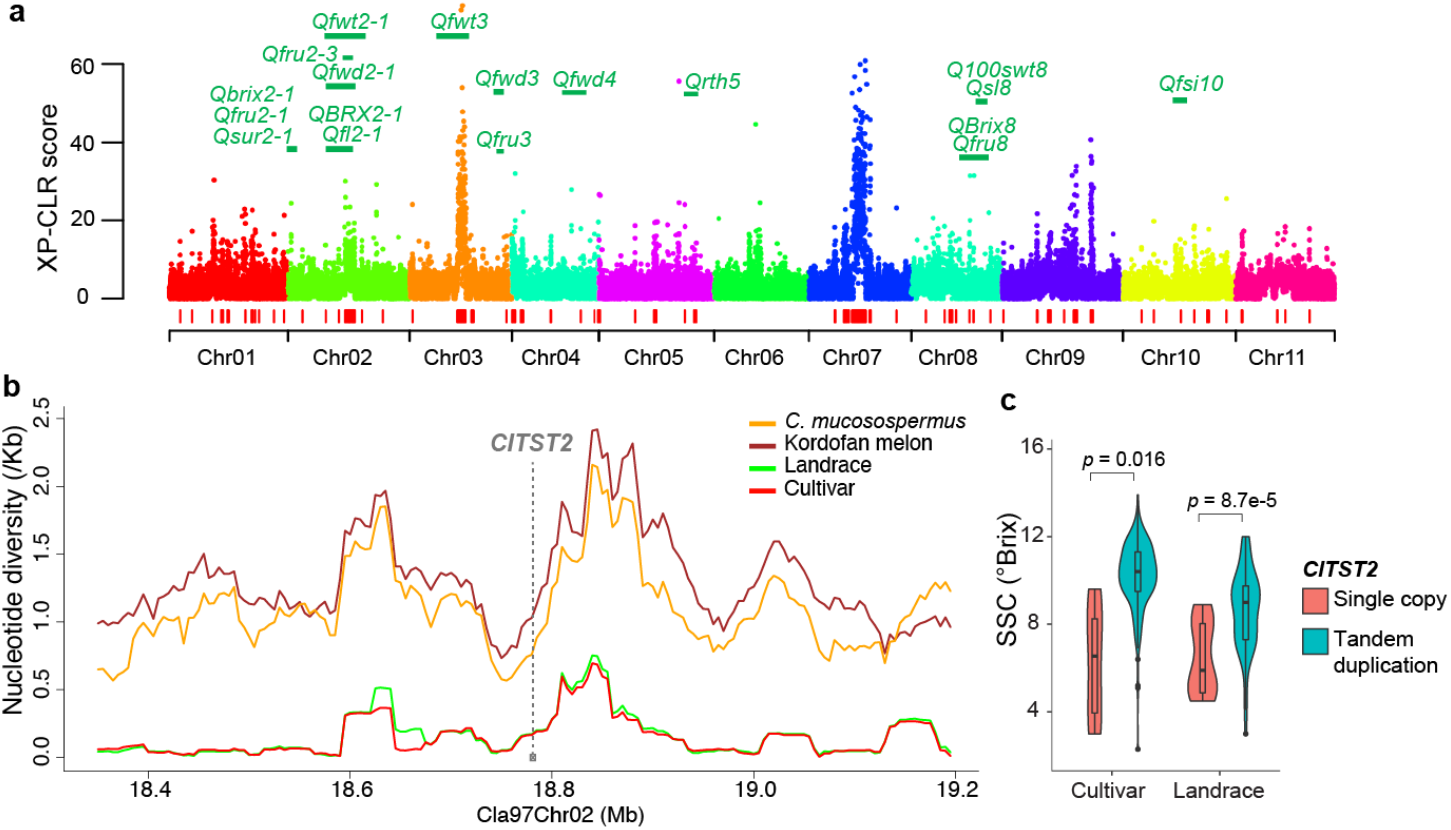
Domestication of cultivated watermelon. (**a**) Domestication sweeps detected through comparing landraces to Kordofan melons. Red bars at the bottom indicate genomic regions under selection. QTLs controlling fruit traits overlapping with domestication sweeps are labeled in green. Fruit flesh sweetness QTLs: *Qbrix2-1, Qfru2-1, Qsur2-1, Qfru2-3, QBRX2-1, Qfru3, QBrix8, Qfru8*; fruit weight QTLs: *Qfwt2-1, Qfwt3*; fruit shape QTLs: *Qfwd2-1, Qfl2-1, Qfwd3, Qfwd4, Qfsi10*; fruit rind thickness QTL: *Qrth5*; seed trait QTLs: *Q100swt8, Qsl8*. (**b**) Nucleotide diversities in genome regions surrounding the *ClTST2* gene in different watermelon populations. (**c**) Comparison of fruit flesh sweetness between accessions carrying the two different alleles of *ClTST2*. SSC, soluble solids content.

The Kordofan melons were not distinctly sweet with flesh soluble solids content (SSC) ranging from 0.2 to 3.2 °Brix (**Supplementary Table 19**). The copy number variation at the sugar transporter *ClTST2* was analyzed in the 547 watermelon accessions based on read alignments at the tandemly duplicated genomic region. Interestingly, among the Kordofan melons only one carried the *ClTST2* tandem duplication, which had the highest SSC (**Supplementary Table 19**). A higher genetic diversity was maintained in the genomic region of *ClTST2* in Kordofan melon compared to landraces (**Fig. 3b**). The *ClTST2* tandem duplication became a predominant allele in landraces (70 out of 86 accessions; 81.4%) and was almost fixed in cultivars (238 out of 245 accessions; 97.1%) (**Supplementary Tables 6** and **20**). Fruit flesh SSC levels were significantly higher in accessions carrying the *ClTST2* tandem duplication compared to the ones with only one copy (**Fig. 3c**), indicating the importance of *ClTST2* copy number variation in controlling fruit flesh sweetness. Accessions of wild watermelon *C. mucosospermus, C. amarus* and *C. colocynthis* mostly carried a single-copy *ClTST2*, with one exception in each species (**Supplementary Table 20**). These results together suggested that the *ClTST2* tandem duplication was present in wild watermelon populations and was selected during domestication likely due to its role in promoting sugar accumulation in fruits.

## Discussion

Wild relatives have been of great interest in disease resistance breeding of watermelon. Here, we generated high-quality reference genomes of wild watermelon species including *C. mucosospermus, C. amarus* and *C. colocynthis* to complement the existing *C. lanatus* references by capturing genomic information unique to these wild watermelons that display tolerance to various biotic and abiotic stresses. Comparative genomic analysis, phylogenetic analysis and molecular dating using these high-quality genomes improved our knowledge on watermelon species divergence and revealed *Citrullus* chromosome karyotype evolution. A *Citrullus* super pan-genome was constructed using reference genome assemblies and genome resequencing data of 547 watermelon accessions spanning four species to represent the watermelon gene repertoire integrated using orthologous relationships across species. Improvement of gene prediction is among one of the most important efforts to build a more complete set of pan-genome genes in a species or genus, alongside with sampling more divergent individuals and generating genome assemblies with high completeness, such as in maize (Hufford et al., 2021). For the *Citrullus* pan-genome, we not only improved gene annotation in different species through homology-based methods to eliminate bias introduced by independent gene predictions, but also established orthologous relationship between predicted genes and genomic locations harboring degenerated copies of ancestral genes to capture gene orthology during evolution and effectively distinguish species-specific genes from divergent orthologs.

The geographic region of watermelon domestication and its direct progenitor have long been under debate, until recently the wild Kordofan melon was identified as the closest relative of the cultivated form (Renner et al., 2021). The Kordofan melons existing in the Darfur region of Northeast Africa are possible relicts of the progenitor population, while the soft-seeded *C. mucosospermus* endemic in West Africa is less likely to be the direct progenitor although it appears to have contributed via introgression (Pérez-Escobar et al., 2022). We compared Kordofan melons with landraces, and identified genomic regions selected during domestication that have not been previously found using *C. mucosospermus* as a representative progenitor population. Combining the information of gene locations within domestication sweeps overlapping with QTLs and gene expression profiles, candidate genes under selection during watermelon domestication were identified for fruit quality traits, such as flesh sweetness, flesh color and rind thickness. In addition, the tandem duplication of the tonoplast sugar transporter gene, *ClTST2*, was found to be associated with increased fruit flesh sweetness in cultivated watermelon. This tandem duplication was present as a rare allele in Kordofan melons and wild watermelon species. It was likely selected during domestication and has become the dominant allele in cultivated watermelon. Red coloration of fruit flesh controlled by *LCYB* was selected during domestication as well. On the other hand, fixation of the loss-of-function allele of *ClBt* in the Kordofan melon population suggests that the non-bitterness trait of fruit flesh could have been fixed before human selection due to genetic drift. Collectively, our *Citrullus* super-pangenome serves as a comprehensive resource for researchers and breeders to mine and utilize genes in cultivated and wild watermelon species. Our comparative analyses using the pan-genome improved our knowledge on gene evolution during watermelon speciation and domestication.

## Methods

### Plant materials and sequencing

Plants of *C. mucosospermus* USVL531-MDR were grown in the greenhouse at Boyce Thompson Institute in Ithaca, New York, with a 16/8 h light/dark cycle at 20°C (night) to 25°C (day). Young leaves from a single 3-week-old plant were collected for high-molecular-weight DNA extraction followed by shearing to fragments with an average size of 20 kb using g-TUBE (Covaris). The sheared DNA was then used to construct a PacBio SMRT library following the standard SMRT bell construction protocol. The library was sequenced on a PacBio Sequel platform using the 2.0 chemistry (PacBio). A paired-end genomic libraries with an insert size of 470 bp was prepared using the Genomic DNA Sample Prep kit (Illumina, San Diego, CA) and sequenced on an Illumina NextSeq 1000 platform. Plants of *C. colocynthis* PI 537277 and *C. amarus* USVL246-FR2 were grown in the greenhouse at the U.S. Vegetable Laboratory, Charleston, South Carolina, with 14-16 h of natural sun light and temperature at 25-30°C. Genomic DNA was extracted from young fresh leaves using the QIAGEN DNeasy Plant Mini Kit (QIAGEN, Valencia, CA) following the manufacturer’s instructions. Paired-end genomic libraries with insert sizes of 200 bp and 500 bp for USVL246-FR2, and 470 bp and 800 bp for PI 537277 were prepared using the Genomic DNA Sample Prep kit (Illumina, San Diego, CA). Four mate-pair libraries with insert sizes of 5, 10 and 15 and 20 kb for USVL246-FR2 and three mate-pair libraries with 2, 5 and 10 kb insert sizes for PI 537277 were prepared. All these paired-end and mate libraries were sequenced on an Illumina HiSeq 1500 system with the paired-end mode. For PI 537277, Hi-C and Chicago libraries were prepared following the protocols implemented by Dovetail Genomics (Scotts Valley, CA, USA) and sequenced on an Illumina HiSeq X platform.

Transcriptome sequencing was performed for samples collected from *C. amarus* USVL246-FR2 leaves and *C. colocynthis* PI 537277 fruit tissues. Total RNA was extracted using QIANGEN RNeasy Plant Mini Kit (QIANGEN). RNA-Seq libraries were constructed using the NEB Next UltraTM RNA Library Prep Kit (NEB, Beverly, MA) and sequenced on a HiSeq 1500 platform.

Fifteen Kordofan melon lines were grown in the greenhouse at Boyce Thompson Institute in Ithaca, New York, with a 16/8 h light/dark cycle and night and day temperature of 20°C and 25°C, respectively. Flowers were hand pollinated at anthesis and fruits were collected at 54 to 74 days after pollination for the measurement of soluble sugar content on a digital refractometer (HR200; APT Co.).

For genome resequencing, young fresh leaf tissue from a single seedling of a total of 202 watermelon accessions including 201 wild accessions and one landrace were collected. Genomic DNA was extracted from the leaf tissue using the Qiagen DNeasy Plant Kit, followed by paired-end library construction using the NEBNext Ultra DNA Library Prep kit according to the manufacturer’s instructions. The libraries were sequenced on an Illumina NextSeq 1000 platform using the paired-end 2 × 150 bp mode.

### *De novo* reference genome assembly

PacBio reads of *C. mucosospermus* USVL531-MDR were error corrected and assembled into contigs using CANU (v1.7.1) (Koren et al. 2017) with default parameters except that both ‘OvlMerThreshold’ and ‘corOutCoverage’ were set to 500. The resulting contigs were then corrected with PacBio long reads using the Arrow program in the SMRT-link-5.1 package (PacBio). Illumina paired-end genome sequencing reads were processed with Trimmomatic (v0.36) (Bolger et al., 2014) to remove adaptors and low-quality sequences. The corrected assembly was subjected to a second-round error correction with the cleaned Illumina reads using Pilon (v1.22) (Walker et al., 2014) with parameters ‘-fix bases -diploid’. Putative contaminations were identified by aligning the assemblies against the NCBI GenBank nucleotide database. Sequences with > 90% of their length aligned to the microbial or organelle genomes were discarded. Redundant contigs that were covered by longer contigs with sequence identity > 99% and coverage > 99% were removed. To construct pseudochromosomes, the contigs were aligned to the 97103 (v2) reference genome using LAST (Kielbasa et al., 2011) to find unique best alignments between USVL531-MDR and 97103. The genetic map constructed using a population derived from a cross between *C. lanatus* cultivar Strain II and *C. mucosospermus* PI 560023 (Sandlin et al., 2012) was used to validate the chromosome-level assembly.

For *C. colocynthis* PI 537277 and *C. amarus* USVL246-FR2, Illumina paired-end and mate-pair reads were used for *de novo* genome assemblies. Trimmomatic (v0.36) (Bolger et al., 2014) and ShortRead (Morgan et al., 2009) were used to remove low-quality and adaptor sequences in reads from the paired-end and mate-pair libraries, respectively. The cleaned reads were assembled into scaffolds with SOAPdenovo2 (Luo et al., 2012), followed by gap filling with the gapcloser program in the SOAPdenovo2 package. Pilon (v1.22) (Walker et al., 2014) was used to correct base errors, fix mis-assemblies and further fill gaps. Contamination and redundant sequences were removed as describe above for USVL531-MDR. For pseudochromosome construction of PI 537277, the scaffolds, cleaned Illumina paired-end reads, Chicago library reads, and Dovetail Hi-C library reads were used as input data for HiRise (Dovetail Genomics), a software pipeline designed for using proximity ligation data to scaffold genome assemblies. The assembly was then manually inspected and curated based on paired-end and mate-pair read alignments. USVL246-FR2 scaffolds were anchored into 11 chromosomes using genetic maps generated from the F2 and RIL populations derived from USVL246-FR2 × USVL114 (Branham et al., 2019).

### Repeat annotation and gene prediction and annotation

Repeat annotation was performed using the EDTA pipeline (Ou et al., 2019). The resulting repeat libraries were used to identify TEs in the genome assemblies with repeatmasker (http://www.repeatmasker.org/). Gene prediction was performed on the repeat-masked genome assemblies with MAKER (Cantarel et al., 2008), which combines evidence from *ab initio* gene prediction, transcript mapping and protein homology to define confident gene models. SNAP (Korf, 2004) and AUGUSTUS (Stanke et al., 2006) were used for *ab initio* gene prediction. *De novo* and genome-guided transcript assemblies were generated with Trinity (Grabherr et al., 2011) using RNA-Seq data generated in this study for *C. amarus* USVL246-FR2 and *C. colocynthis* PI 537277, and previously published 97103 RNA-Seq data (NCBI SRA accession SRP012849) for *C. mucosospermus* USVL531-MDR. The two types of transcript assemblies were then combined and aligned to the genomes using the PASA2 pipeline (Haas et al., 2003). The resulting alignments were used as the transcript evidence. To provide protein homology evidence, protein sequences from Arabidopsis (TAIR 10), watermelon (97103 v2), cucumber (Gy14 v2), and melon (DHL92 v4.0), as well as the UniProt (Swiss-Prot plant division) database were aligned to the genomes using Spaln (Iwata and Gotoh, 2012). Furthermore, gene predictions of the six watermelon reference genomes, including three developed in this study and three published ones (Guo et al., 2019; Wu et al., 2019; Renner et al., 2021), were improved through mapping genes between assemblies with Liftoff (Shumate and Salzberg, 2021). Briefly, coding sequences (CDS) of genes predicted in the other five watermelon reference genomes were aligned to one target genome, requiring one-to-one mapping with at least 90% identity and 90% coverage. Gene models originally predicted in the target genome and those ‘lifted over’ from the other five genomes were integrated, with the longest gene model kept for each gene. When adding a new gene model to a genomic region based on the Liftoff results, only non-TE genes containing InterPro domains were considered.

For gene annotation, protein sequences of the predicted genes were compared against the GenBank nr, the UniProt (Swiss-Prot and TrEMBL; http://www.uniprot.org/) and Arabidopsis (TAIR 10) databases using BLAST (Camacho et al., 2009) with an E-value cut-off of 1e-4, as well as the InterPro database using InterProScan (Jones et al., 2014). Gene ontology (GO) annotations were obtained using Blast2GO (Conesa et al., 2005) based on the BLAST results against nr and the InterProScan analysis. BLAST results against UniProt and TAIR were processed using AHRD (https://github.com/groupschoof/AHRD) for assigning functional descriptions to the predicted genes.

### Divergence time and genome evolution

To estimate the divergence times among *Citrullus* species, genes in the genomes of five watermelons (*Citrullus lanatus* subsp. *vulgaris, C. lanatus* subsp. *cordophanus, C. mucosospermus, C. amarus*, and *C. colocynthis*), seven other cucurbit species (bottle gourd, bitter gourd, cucumber, melon, monk fruit, snake gourd, and sponge gourd), walnut (Zhang et al., 2020) and woodland strawberry (Edger et al., 2018) were used to perform molecular dating. Gene sequences of the cucurbit species were downloaded from GuGenDBv2 (Yu et al., 2023). CDS sequences of single-copy orthologous genes identified by OrthoFinder (v 2.5.4) (Emms and Kelly, 2019) were concatenated and aligned using MUSCLE (v5) (Edgar, 2004). Bayesian divergence time estimation was performed using the MCMCtree program in PAML (v 4.10.5) (Yang, 2007) with the known divergence time between walnut and Cucurbitaceae (84-105 Mya) as the calibration node, 20,000 burn-in iterations, 500 sampling frequency, and 20,000 accepted samples. Chromosome synteny comparison was performed with MCScan (https://github.com/tanghaibao/jcvi/wiki/MCscan-(Python-version)) using gene sequences of melon (DHL92 v4.0), *C. colocynthis* PI 537377, *C. amarus* USVL246-FR2, *C. mucosospermus* USVL531-MDR and *C. lanatus* 97103.

### Construction and annotation of watermelon species-level pan-genomes

Raw Illumina reads were processed to trim adapters and low-quality sequences using Trimmomatic (v0.36) (Bolger et al., 2014). The high-quality cleaned Illumina reads from each accession were *de novo* assembled using SPAdes (v3.12.0) (Bankevich et al., 2012) with parameters ‘--careful -k 55,77,99,127’. Assembled contigs with length ≥500 bp were kept. The assembled contigs were aligned to the reference genome of the same species using QUAST (v5.0.2) (Gurevich et al., 2013) with parameters ‘--min-identity 90.0 -- min-alignment 300’. Genomes of 97103, USVL531-MDR, USVL246-FR2, and PI 537277 were used as the references for species *C. lanatus, C. mucosospermus, C. amarus*, and *C. colocynthis*, respectively. Unaligned sequences with length ≥500 bp were extracted and searched against the NCBI GenBank nucleotide database using BLAST (Camacho et al., 2009) to identify possible contaminations. Sequences with best hits from outside the green plants, or covered by known plant chloroplast or mitochondrial genomes, were removed. The cleaned non-reference sequences from all accessions of the species were combined and consolidated into unique sequences using CD-HIT (Li and Godzik, 2006) with an identity threshold of 0.9. Redundancy was further removed by performing all-versus-all alignments using BLAST with an identity threshold of 90%. The resulting cleaned non-redundant non-reference sequences and the reference sequences were merged as the species-level pan-genome. Protein-coding genes were predicted from non-reference sequences as described above for the reference genome assemblies.

### Construction of the *Citrullus* super-pangenome

To build the *Citrullus* super pan-genome, orthologous relationships were established among the species-level pan-genome genes. CDS of genes in one species-level pan-genome were mapped to the other three pan-genomes to determine the orthologous genes or genomic regions using Liftoff (Shumate and Salzberg, 2021), requiring one-to-one alignment with both identity and coverage ≥90%. Gene pairs with at least 50% overlapped CDS regions were considered as orthologous genes. Otherwise, the mapped genomic locations were recorded as the orthologous regions. The gene-to-gene and gene-to-location orthologous relationships identified from the pairwise bidirectional gene ‘lifting over’ were consolidated into orthologous clusters containing the four species. Singleton genes that could be aligned to the sequences in the orthologous clusters using Blat (Kent, 2002) with at least 95% of identity and 50% of coverage were further added to the existing orthologous groups. Remaining singletons were aligned to the pan-genomes using Blat (Kent, 2002), requiring a minimal identity of 95% and a minimal coverage of 90% to identify additional gene-to-location relationships. Syntenic genomic regions across species were determined based on the syntenic gene blocks identified using MCScanX (Wang et al., 2012) with gene anchors whose CDS sequences were reciprocal best hits between species. Based on the syntenic genomic regions, the original orthologous groups were then categorized into syntenic orthologous groups and orthologous groups without syntenic information.

### Gene PAV analysis

Genomic reads from each accession were aligned to the pan-genome of its own species using BWA-MEM (v0.7.17) (Li, 2013) with default parameters. The read coverage of the pan-genome was then computed from the alignments using the ‘genomecov’ utility in the BEDTools suite (Quinlan and Hall, 2010) with parameters ‘-bg -split’. To control false negative calling of gene PAVs, accessions with sequences covering less than 320 Mb of the pan-genome with at least one read were excluded from the gene PAV analysis. Read coverages for genes were calculated using BEDTools ‘intersect’ with the parameter ‘-wao’ based on gene positions. A gene with 50% of its CDS length covered by one read was considered as present in the accession. Genes with significantly changed occurrence frequencies between two groups were determined using the Fisher’s exact test with FDR corrected for multiple comparisons.

For genotyping at *ClTST2* in watermelon accessions, a junction site uniquely present in the tandem duplication allele (in the 97103 reference) was identified. Illumina paired-end reads were aligned to the 97103 reference using BWA-ALN (v0.7.17) (Li, 2013) with parameters ‘-n 0.01 -o 1 -e 2’ and ‘bwa sampe -s’. Read alignments supporting the presence of the junction site were used to indicate the presence of the *ClTST2* tandem duplication (**Supplementary Fig. 12**). Read counts were also calculated for a nearby genomic site (500 bp upstream of the junction site) as a control. Read alignments were found at the control site for all accessions in this study, and the lack of read support at the junction site was considered absence of tandem duplication or single-copy *ClTST2*.

### Variant calling, phylogenetic analysis and selective sweep identification

Raw Illumina DNA reads were processed using Trimmomatic (v0.36) (Bolger et al., 2014) to remove low quality and adaptor sequences. The cleaned reads were aligned to the 97103 genome using BWA-MEM (v0.7.17) (Li, 2013) with default parameters. SNP and small indel calling was performed using the Sentieon software package (https://www.sentieon.com/). Briefly, duplicated read pairs in each alignment file were marked using the Sentieon Dedup function, and variants from each sample were then called with the Sentieon Haplotyper function, followed by joint variant calling using Sentieon GVCFtyper. Hard filtering was applied to the raw variant set using GATK (v4.1) (McKenna et al., 2010), with parameters ‘QD < 2.0 || FS > 60.0 || MQ < 40.0 || MQRankSum < -12.5 || ReadPosRankSum < -8.0’ for SNPs and ‘QD<2.0 || FS>200.0 || ReadPosRankSum <−20.0’ for small indels. Only bi-allelic variants with minor allele frequency (MAF) > 0.01 were used in the downstream analyses. Variants were annotated using SnpEff (v5.0e) (Cingolani et al., 2012).

Phylogenetic relationships among the 547 watermelon accessions were inferred using 106,430 SNPs at the fourfold degenerate sites. A maximum likelihood tree was constructed using IQ-TREE (v1.6.12) (Nguyen et al., 2015) with 1,000 bootstrap replicates and a *C. naudinianus* accession, PI 596694, as the outgroup. Cross-population composite likelihood ratio test was performed using XP-CLR (v1.0) (Chen et al., 2010) to scan the 97103 genome for domestication sweeps by comparing landraces to Kordofan melons with parameters ‘-w1 0.0005 100 100 1 -p0 0.7’. Genetic distances between adjacent SNPs were calculated based on physical distance of adjacent markers in an integrated genetic map (Ren et al., 2014). Genomic regions with XP-CLR scores in the top 10% overlapping with top 50% of π ratios of Kordofan melons to landraces were identified as potential selective sweeps.

## Data availability

This Whole Genome assemblies been deposited at DDBJ/ENA/GenBank under the accessions JAKCFO000000000, JALMEV000000000 and QLSR00000000 for *C. mucosospermus* USVL531-MDR, *C. amarus* USVL246-FR2, and *C. colocynthis* PI 537277, respectively. Raw genome and transcriptome sequencing reads have been deposited in the NCBI BioProject database under the accession numbers PRJNA476359, PRJNA808410 and PRJNA794184. Genome (http://cucurbitgenomics.org/v2/genome_summary) and pan-genome (http://cucurbitgenomics.org/v2/ftp/pan-genome/watermelon/super-pangenome/) assemblies and annotation, and SNPs and small indels in VCF file format (http://cucurbitgenomics.org/v2/ftp/reseq/watermelon/v3/) are also available at CuGenDBv2 (Yu et al., 2023).

### Acknowledgements

This research was supported by grants from USDA National Institute of Food and Agriculture Specialty Crop Research Initiative (2015-51181-24285 and 2020-51181-32139) and the US National Science Foundation (IOS-1855585).

## Author contributions

Z.F. and S.W. designed and managed the project. S.B., W.P.W., C.K., A.L., C.M., S.S.R., X.Y. and Z.F. provided plant materials and/or contributed to DNA and RNA sequencing. S.H, S.W. and L.G. performed data analyses. S.W. and H.S. wrote the manuscript. Z.F. revised the manuscript.

## Competing interests

The authors declare no competing interests.

## Notes

### Competing Interest Statement

The authors have declared no competing interest.

### Summary of Updates

Add a co-author

