## Supplementary figures for "A *Citrullus* genus super-pangenome reveals extensive variations in wild and cultivated watermelons and sheds light on watermelon evolution and domestication"

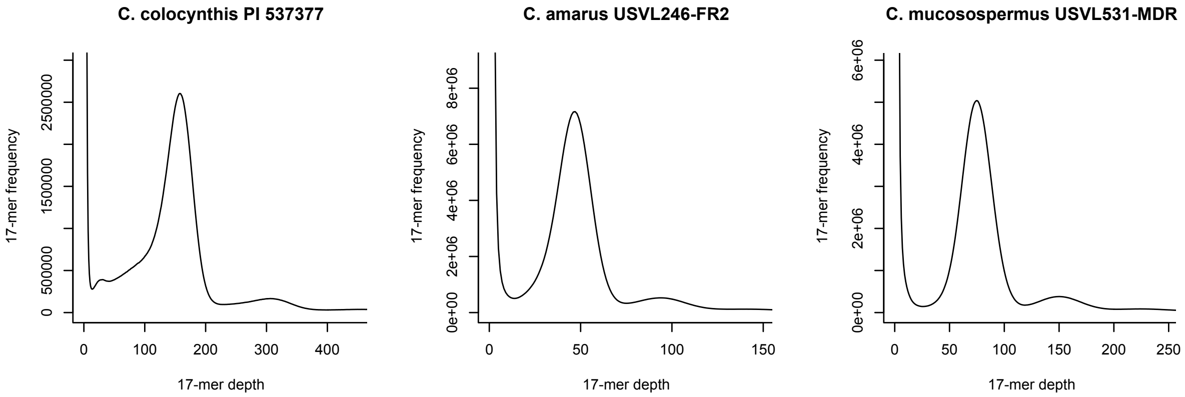


**Figure S1** K-mer distribution of Illumina genomic sequencing reads of *C. colocynthis* PI 537277, *C. amarus* USVL246-FR2, and *C. mucosospermus* USVL531-MDR. Genome sizes of USVL531-MDR, USVL246-FR2 and PI 537277 were estimated to be 434.7 Mb, 423.2 Mb and 406.0 Mb, respectively. K-mer (k=17) counting was performed with Jellyfish (https://github.com/gmarcais/Jellyfish) and the genome size was estimated based on the formula: Genome size = total number of k-mers / peak depth.

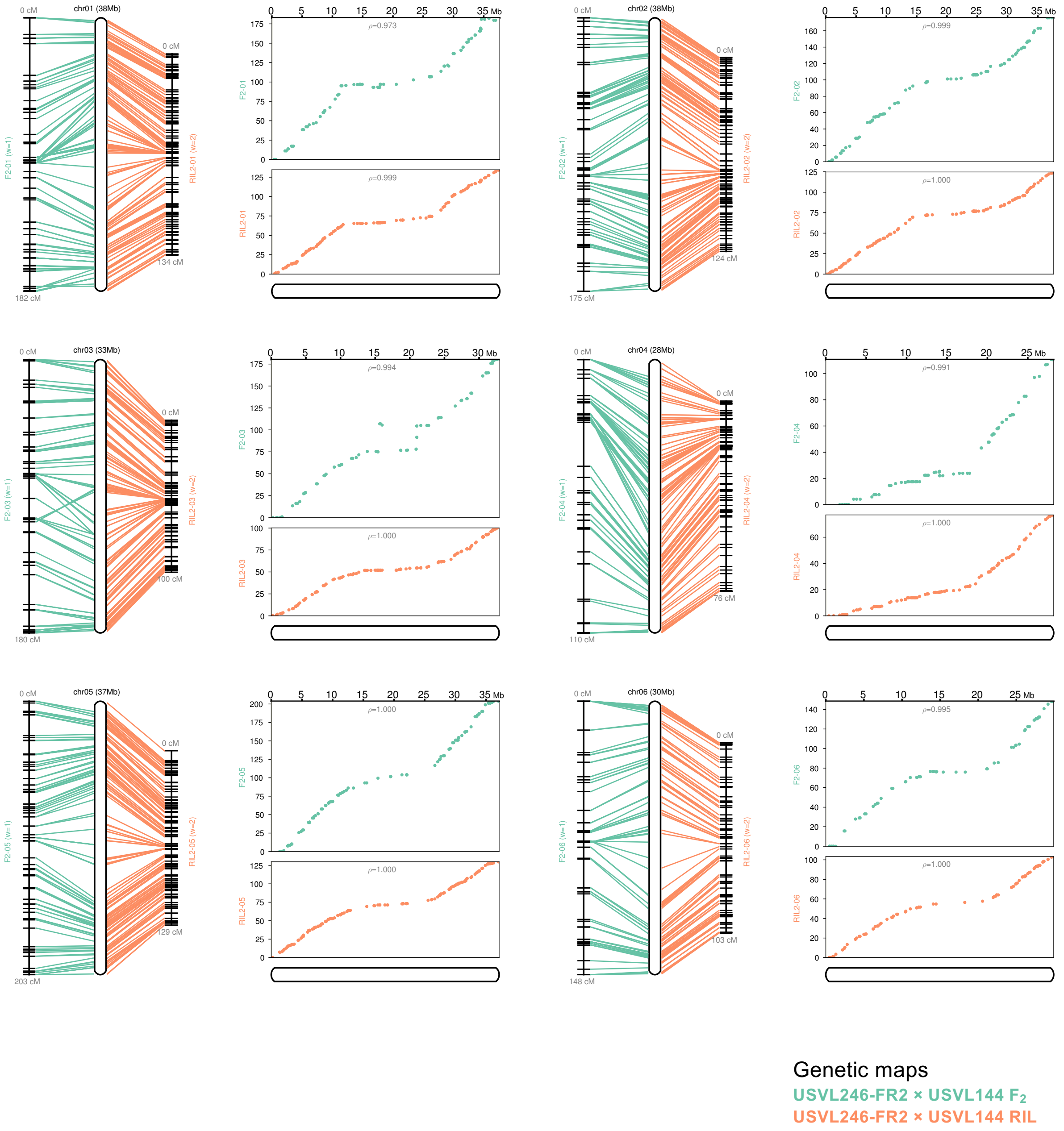


**Figure S2** Collinearity between the USVL246-FR2 pseudomolecules and genetic maps.

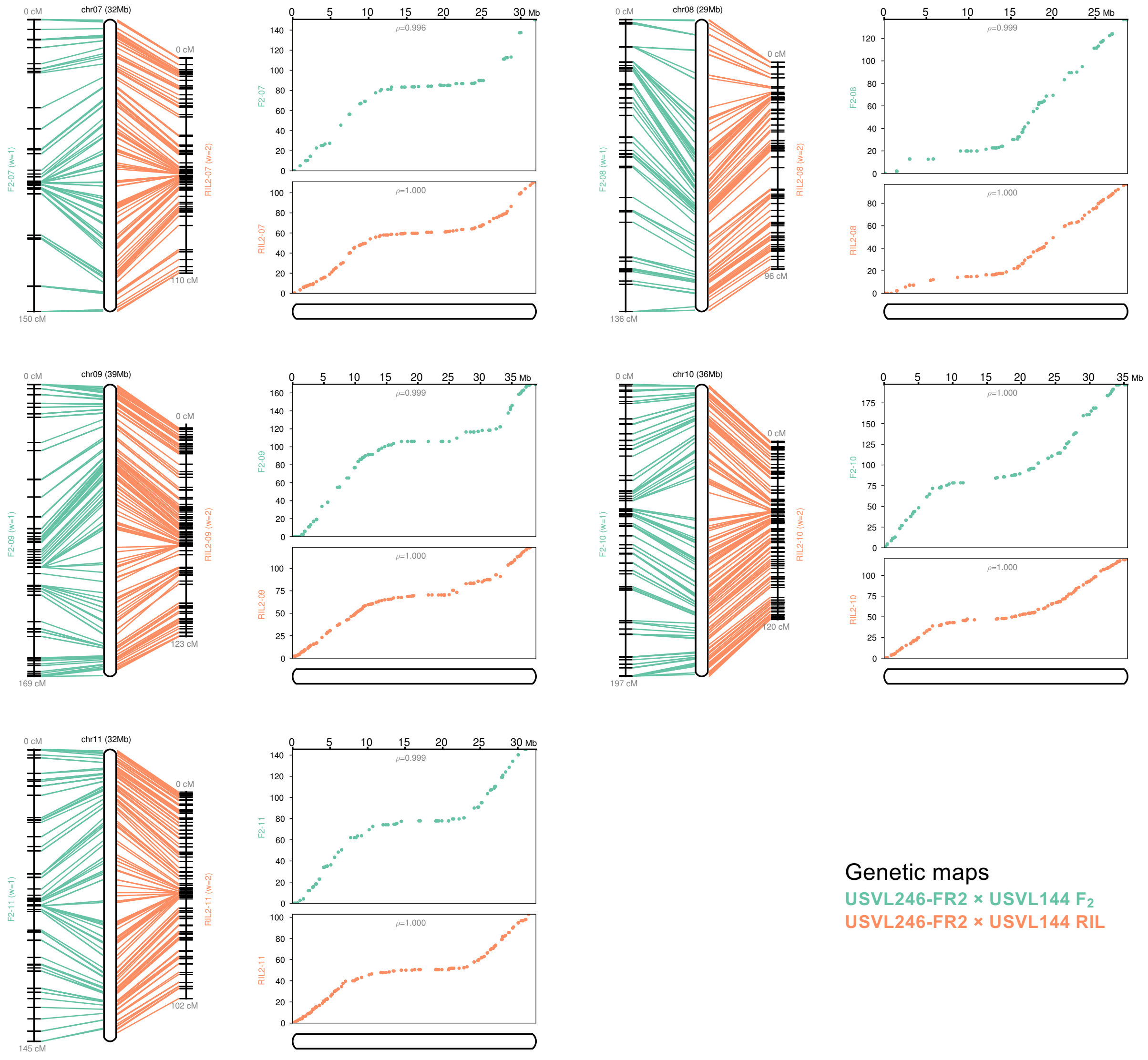


**Figure S2** Continued

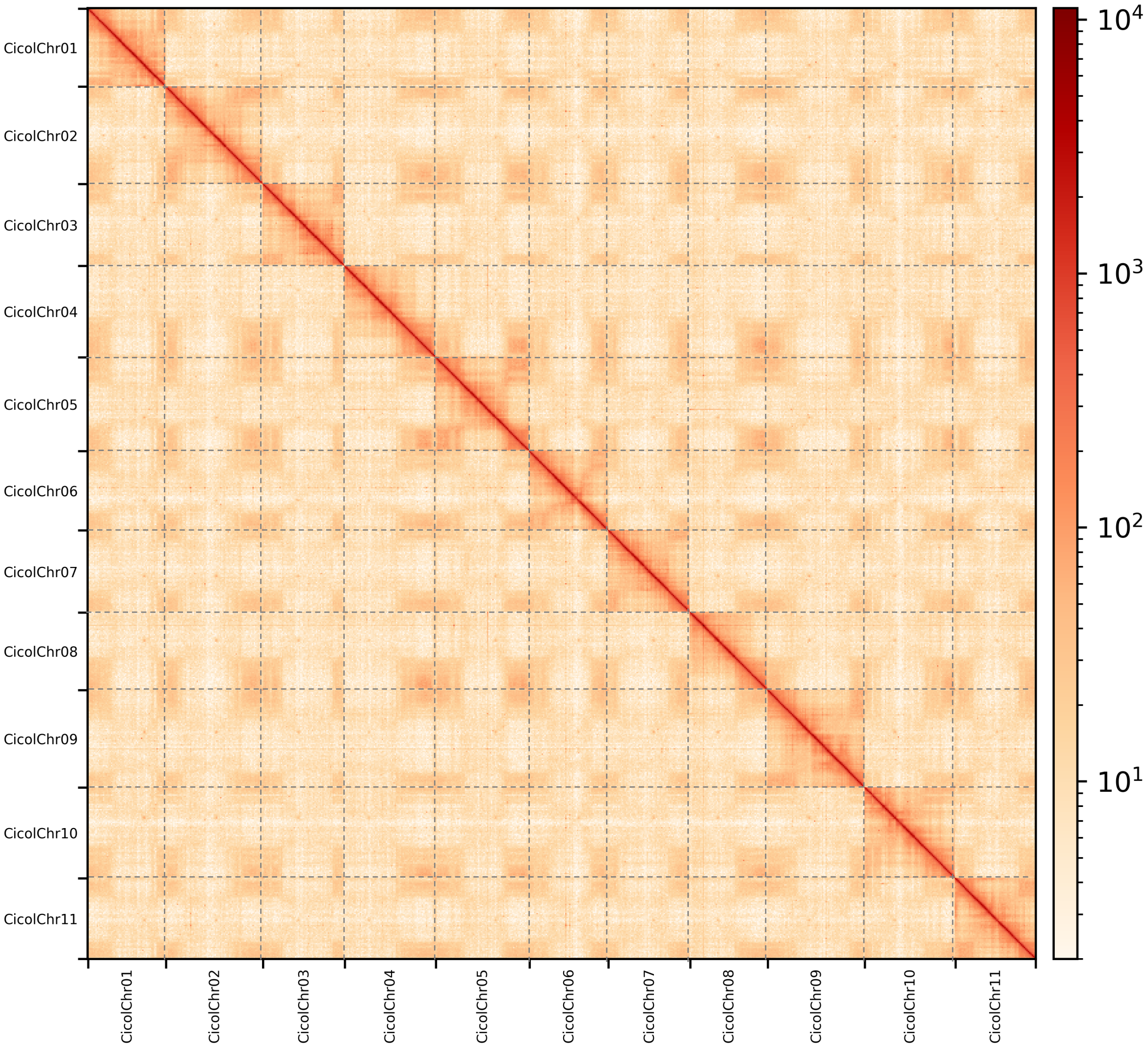


**Figure S3** Hi-C chromatin interaction heatmap of *C. colocynthis* PI 537277.

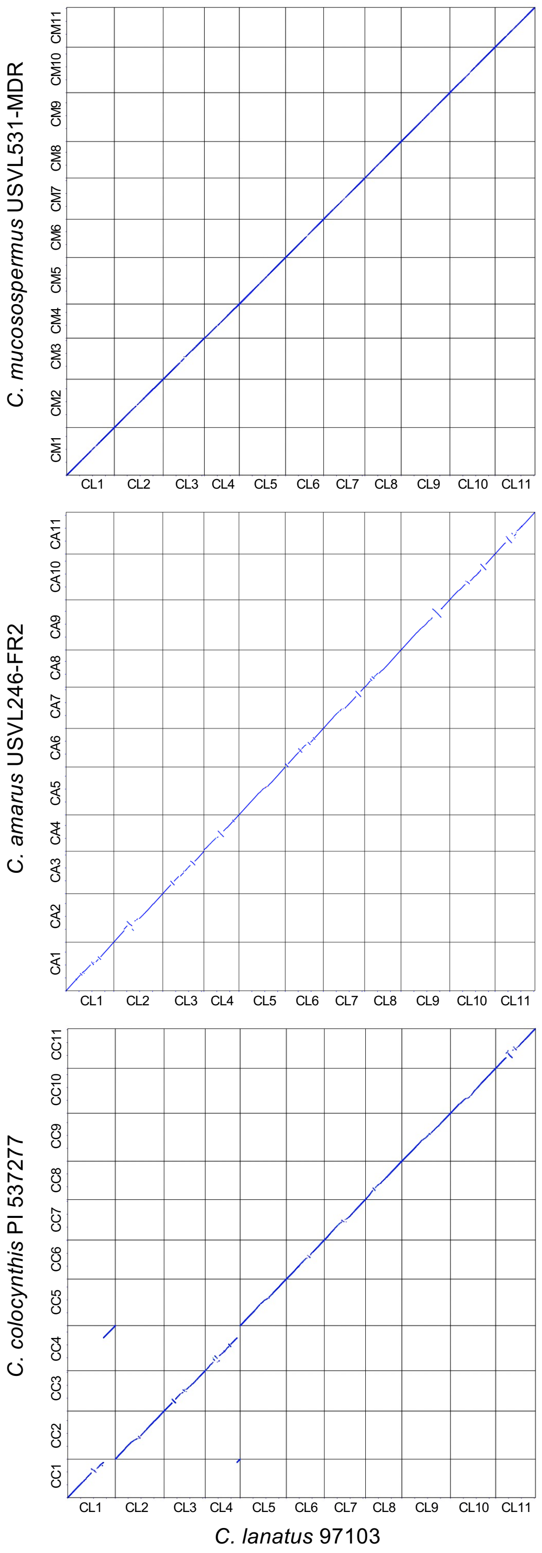


**Figure S4** Collinearity between genomes of the cultivated watermelon 97103 and wild watermelons *C. mucosospermus* (CM) USVL531-MDR, *C. amarus* (CA) USVL246-FR2, and *C. colocynthis* (CC) PI 537277.

(a)

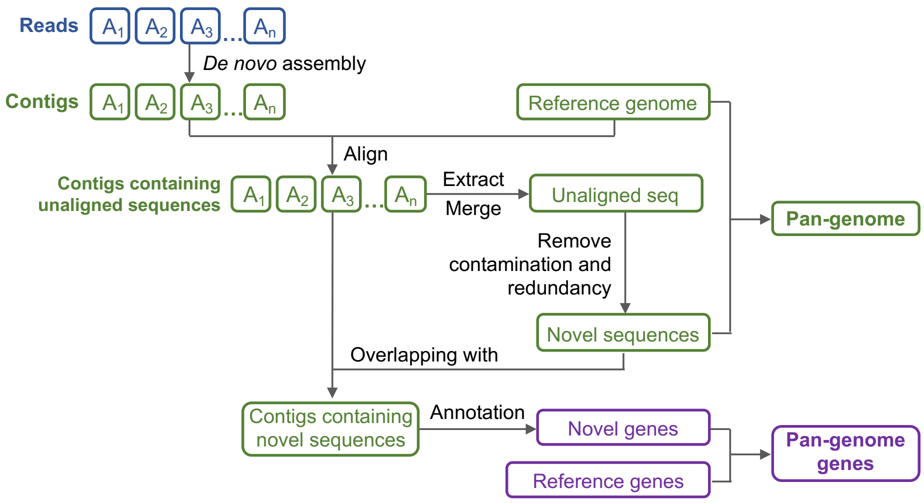


(b)

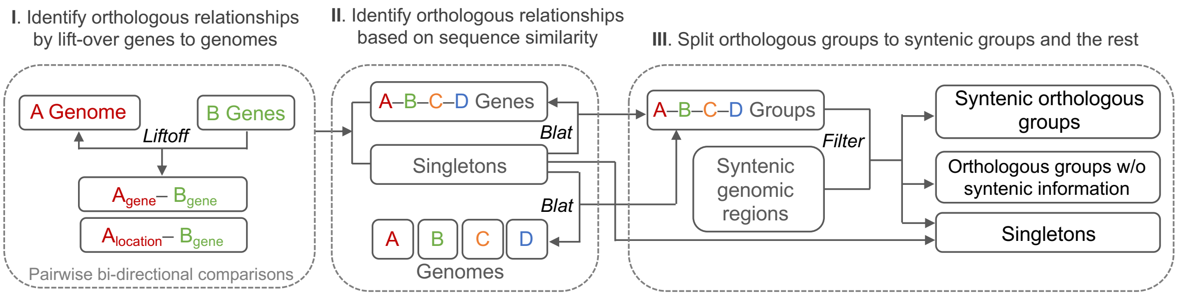


**Figure S5** Workflow for the construction of the *Citrullus* super-pangenome. (a) Strategy for construction of species-level pan-genome by aligning *de novo* assembled sequences of individual accessions (A1, A2, A3, …, An) to the species-specific reference genome followed by identification of non-redundant novel sequences. (b) Strategy for construction of the *Citrullus* super-pangenome through establishing gene-to-gene and gene-to-location orthologous relationship among species-level pan-genomes. The colored A, B, C and D letters represent four *Citrullus* species.

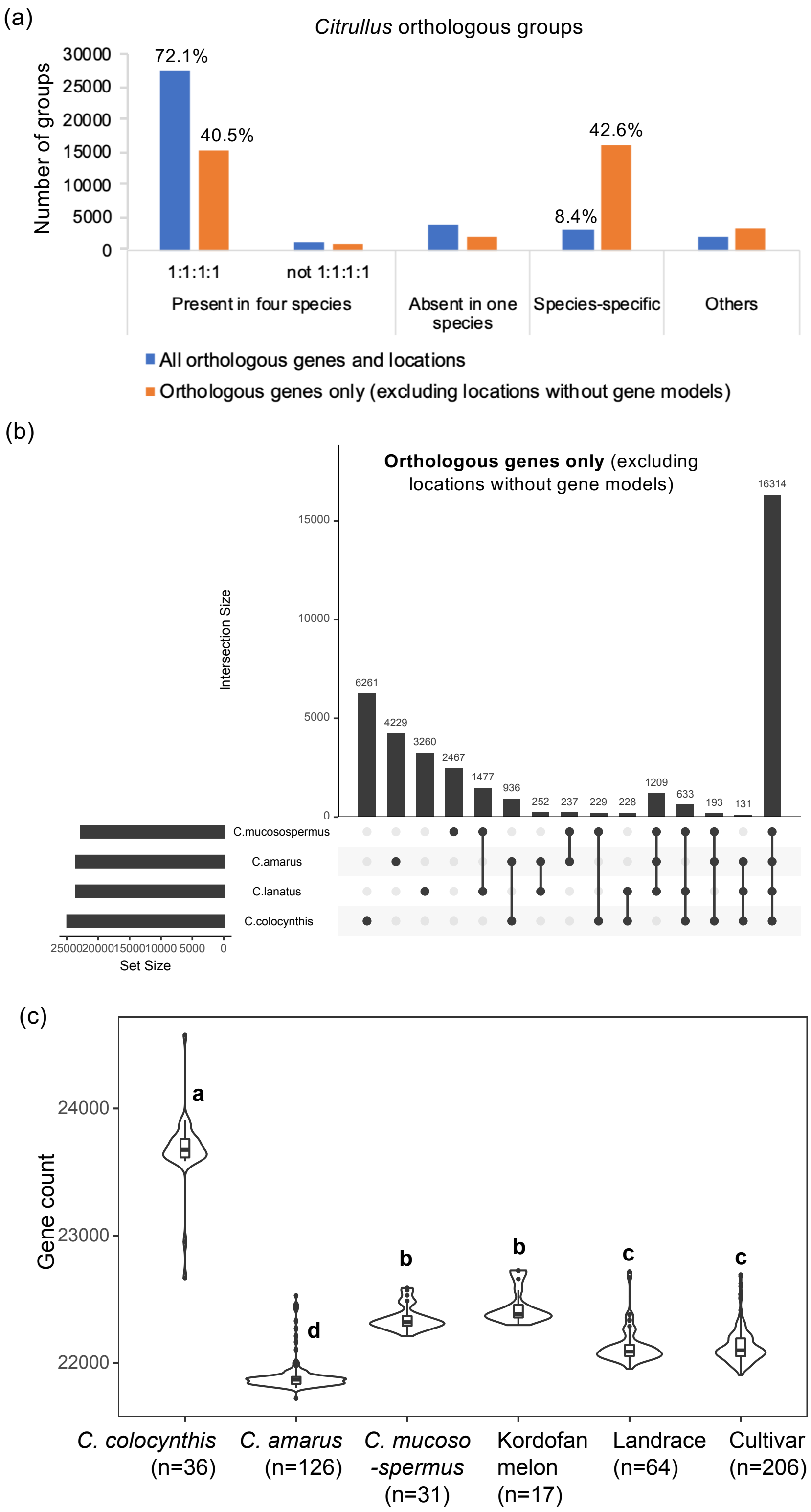


**Figure S6** Genes in the *Citrullus* super-pangenome. (a) Proportions of shared and species-specific orthologous groups. (b) Upset diagram of orthologous gene groups among the four watermelon species, with orthologous locations without predicted genes excluded. (**c**) Numbers genes (predicted gene models only; excluding orthologous locations without predicted genes) detected in individuals of different watermelon populations. Different letters in the violin plots indicate the significant differences among different groups evaluated by Tukey'sHSD test (α<0.05).

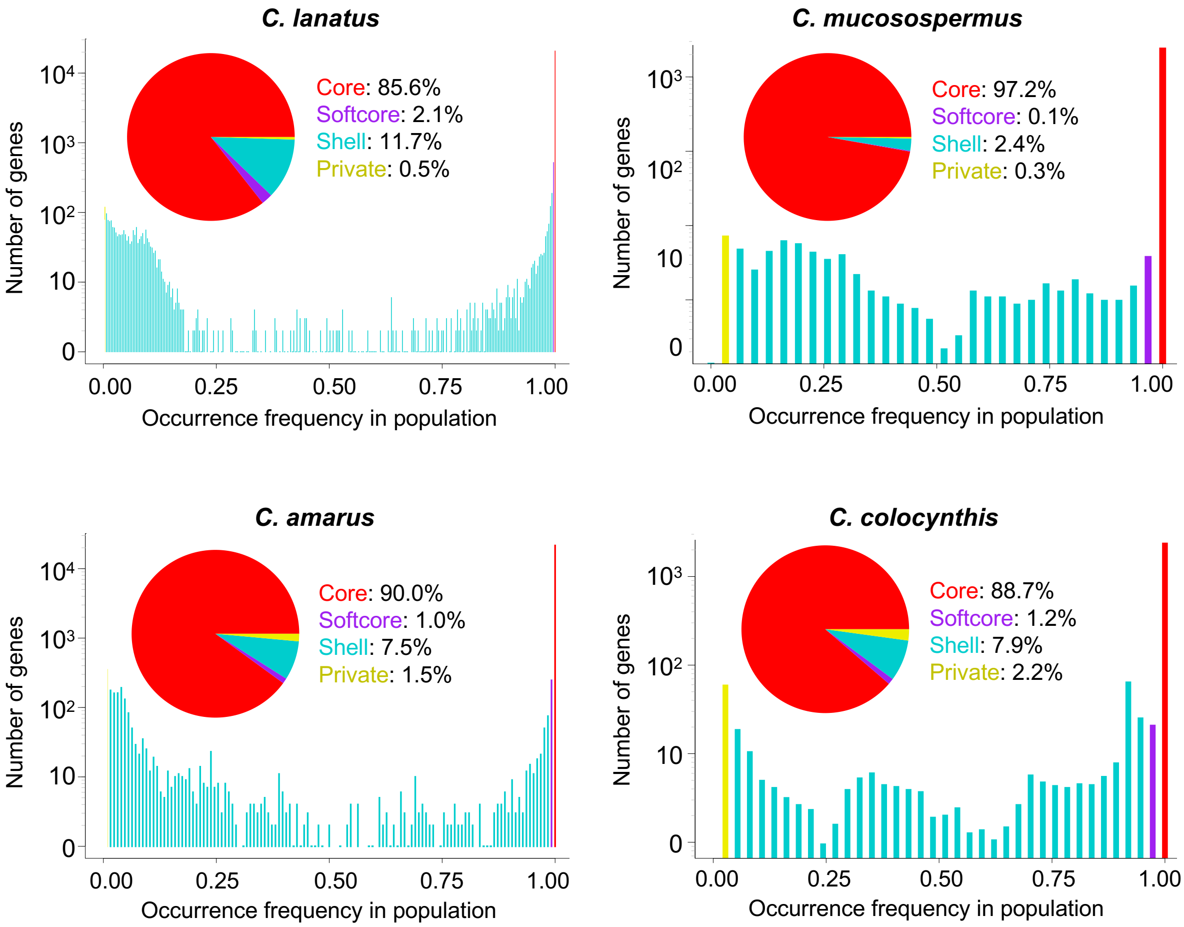


**Figure S7** Compositions of the four species-specific pan-genomes. Core: present in all accessions; softcore: present in all but one; private: present in only one accession; shell: between softcore and private.

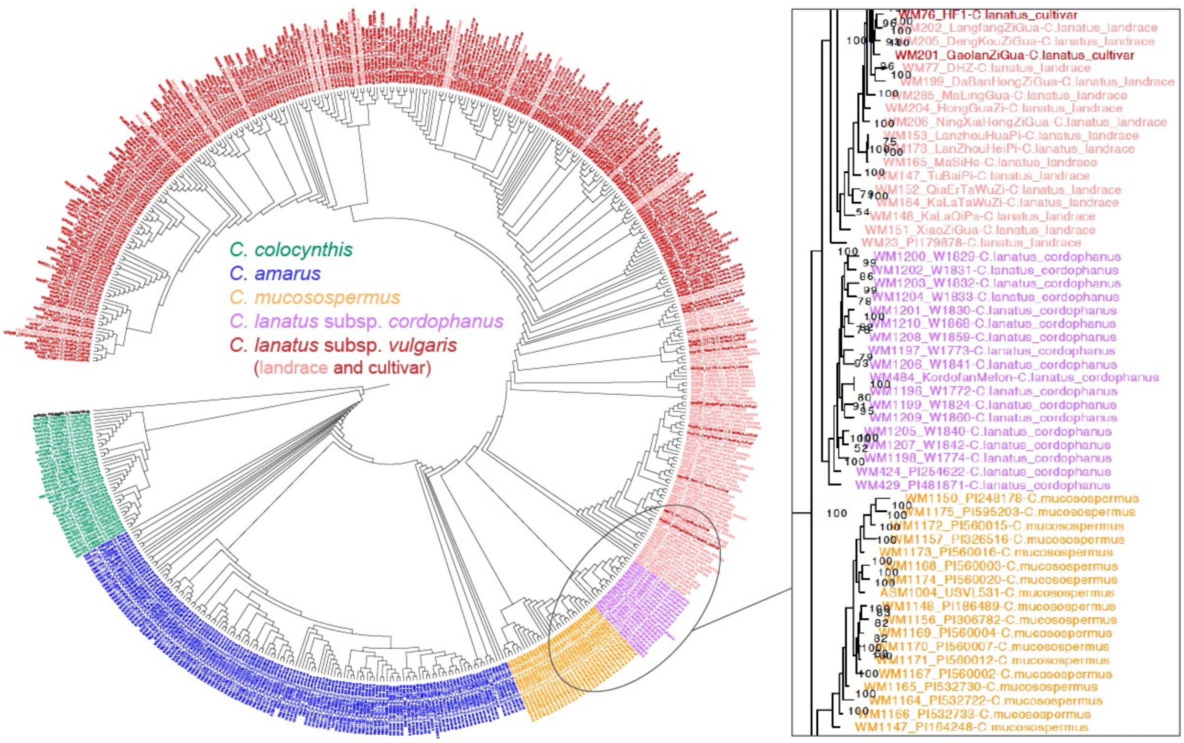


**Figure S8** Maximum likelihood phylogenetic tree of wild and cultivated accessions.

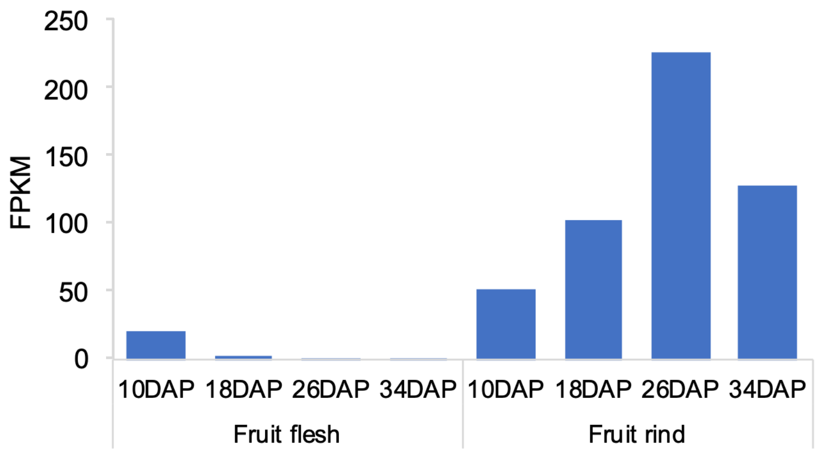


**Figure S9** Expression levels of *Cla97C05G101010* in watermelon 97103 fruit flesh tissues at different developmental stages. DAP, days after pollination.

**a**

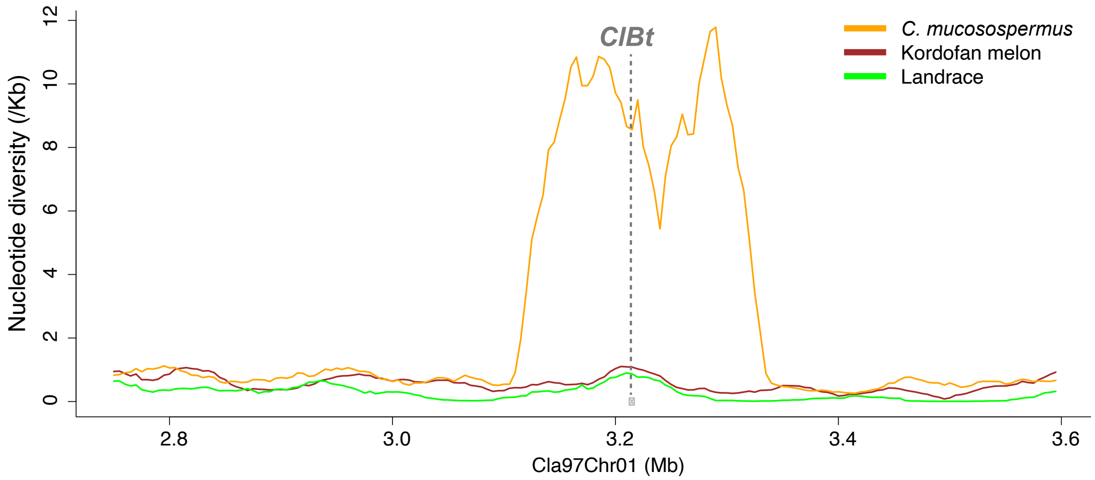


**b**

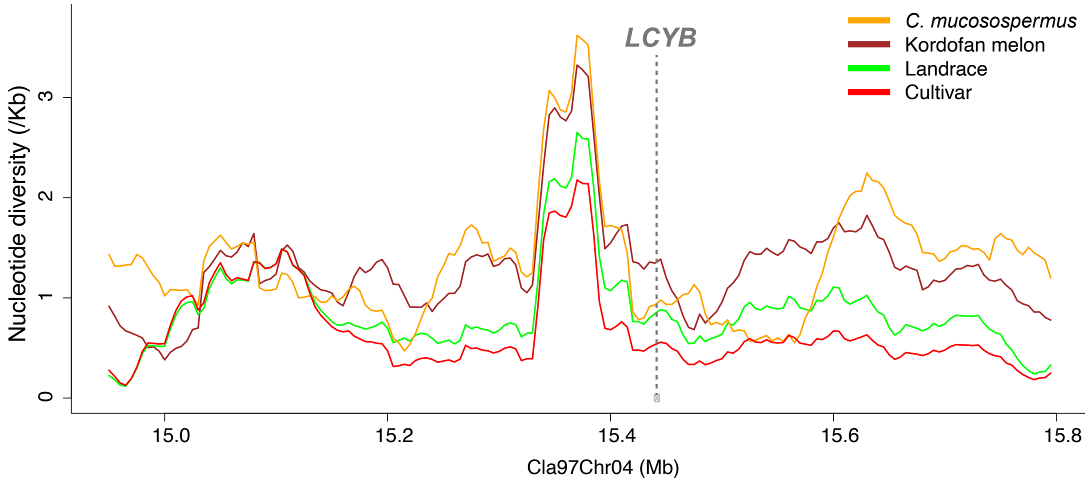


**Figure S10** Nucleotide diversities in the genomic regions surrounding the *ClBt* (**a**) and *LCYB* (**b**) genes in different watermelon populations.

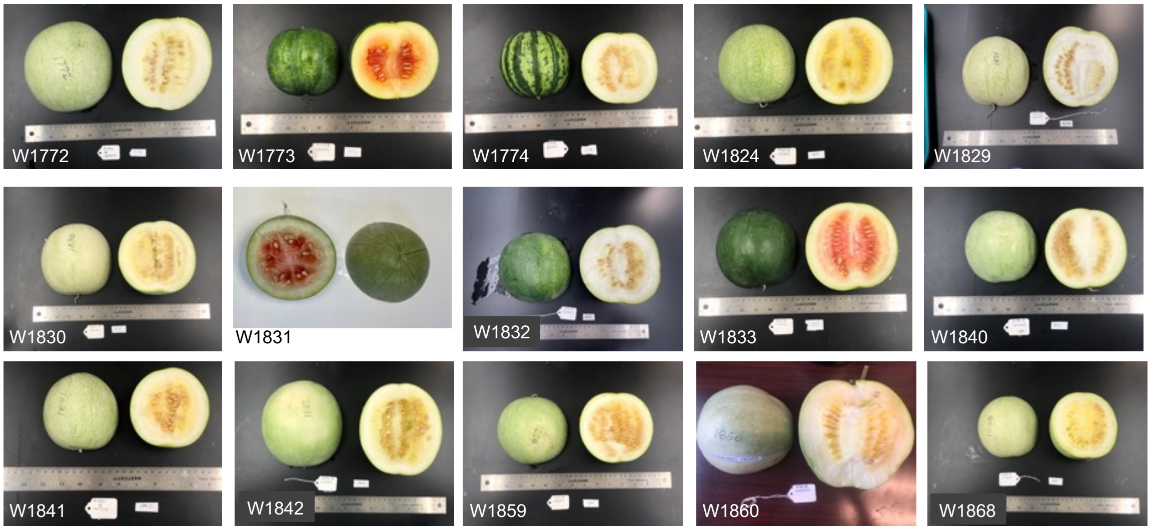


**Figure S11** Mature fruits of 15 Kordofan melon accessions.

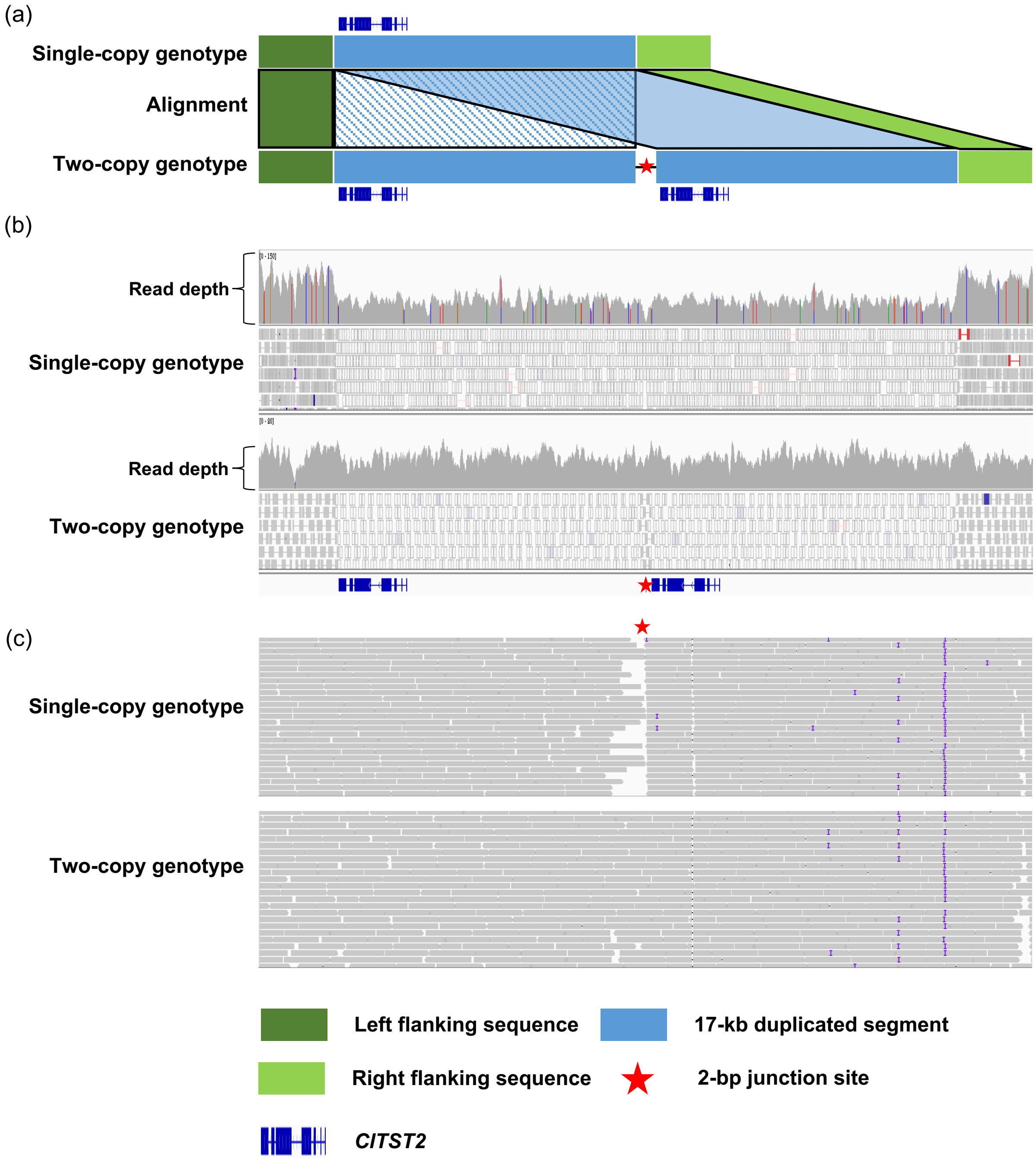


**Figure S12** Read alignment at *ClTST2*. (a), diagram of the *ClTST2* tandem duplication. (b), read depth in the genomic region harboring the *ClTST2* tandem duplication. The accession with single-copy *ClTST2* has reduced read depth (about half) compared to the flanking regions (top panel), while the accession with two copies has similar length to the flanking regions (bottom panel). (c), read alignment at the 2-bp junction site unique to the *ClTST2* tandem duplication allele. Reads generated from the accessions with single-copy *ClTST2* cannot span the junction site (top panel), while those from the accession with two copies can (bottom panel).
